# The disadvantage of having a big mouth: the relationship between insect body size and microplastic ingestion

**DOI:** 10.1101/2025.01.16.633357

**Authors:** M.W. Ritchie, E.R. McColville, J.E. Mills, J.F. Provencher, S.M. Bertram, H.A. MacMillan

## Abstract

Plastic pollution is ubiquitous, and animals are exposed to diverse plastic shapes and sizes. When plastics enter natural environments, they break down into microplastics (MPs; <5 mm) and likely become more accessible to smaller animals. Insects play critical environmental and economic roles, ingest plastics in the wild, and can physically degrade ingested MPs into smaller and more harmful nanoplastics. While particle size and body size undoubtedly impact plastic ingestion, we have no predictive understanding of how these factors interact to influence which plastics are a threat to which animals. To uncover these potential interactions, we studied how a model cricket species (*Gryllodes sigillatus*) interacts with plastics of differing sizes throughout a twentyfold change in body size during growth and development. We fed crickets a range of MP sizes of 38 to 500 µm with clearly defined particle size thresholds. We investigated whether crickets would avoid MPs when given a choice and found that they do not; instead they gradually began to consume more of the plastic diet over time. We then studied how MP ingestion is influenced by body size and mouth size, and the extent of breakdown that occurs once MPs are ingested. We found that crickets would only consume whole beads when their mouth size was larger than the MP. While small MPs were more likely to be excreted whole, larger MPs were more extensively broken down as crickets grew. We conclude that crickets do not exhibit avoidance behaviour towards plastic and ingest it once a particle can be consumed whole. These effects of insect behaviour and body size on the likelihood of plastic ingestion and the degree to which MPs are degraded have important implications for regulating the size classes of plastic particles entering natural environments and how plastics move through those environments once discarded.

## Introduction

Plastic is an integral part of modern life because of its strength, low weight, and low production costs, but its extensive use has led to significant environmental challenges. Large plastics entering natural environments break down over time into smaller pieces due to abiotic factors such as solar radiation and tidal currents (Zantis et al., 2021; K. Zhang et al., 2021). Eventually, these large plastics break down into sizes between 5 mm and 1 µm, a size class known as microplastics (MPs) (Li et al., 2024; Thompson et al., 2004).

The 5000-fold range in sizes among MPs undoubtedly leads to variation in the availability of plastics to different organisms, and different sizes of plastics may have widely varying consequences for the organisms that ingest them. A diverse suite of MPs composed of different polymers are present in marine and terrestrial environments, where they are ingested by animals, including aquatic mammals, fish and invertebrates (Daghighi et al., 2023; Park et al., 2022; Zantis et al., 2021). Terrestrial insects also encounter MPs in their environments, particularly those living in agricultural settings where sludge from wastewater treatment (biosolids) is being actively applied as fertilizer (Corradini et al., 2019; McColville et al., 2023; Rillig, 2012). The MPs within these biosolids exist in various non-uniform sizes and morphologies (Sivarajah et al., 2023).

The wide variety of MPs presents challenges in determining whether particles ranging from 1 µm to 5 mm differentially affect insects. Additionally, the enormous variety in animal body size in general (e.g. million-fold differences in body weight within animal taxa (Lindstedt & Hoppeler, 2023; M. Silva & Downing, 1995) and insect body mass (six magnitude change in Order (Dillon & Frazier, 2013)) emphasizes the importance of studying MP interactions and fitness effects using particle sizes relevant to a species of interest. For example, an upper limit to the size of ingestible MPs of approximately 125 µm has been found for *Chironomus riparius* larvae (a freshwater non-biting midge) (Silva et al., 2019). For the same species, smaller particles ranging from 1-90 µm were only ingested once the head capsule width of *C. riparius* grew (Scherer et al., 2017). The authors proposed that the structural limitations of the mouth might influence the size of microplastics that can be ingested, as well as the maximum size that can be consumed. Beyond morphological effects, what sizes of MPs are ingested may also depend on the behaviours insects exhibit when encountering plastics in their natural habitats, which may not be consistent among species or over time. For instance, ants have been observed to lose their aversion to food contaminated with 10 µm plastic over time, and their ability to distinguish between contaminated and uncontaminated food diminishes (Le Hen et al., 2024).

The size of ingested plastics matters for insects; several accounts have shown that the size of the ingested MPs can greatly affect fitness-related traits in invertebrates. Large MPs can harm the microbiome of mealworms (Brandon et al., 2018). Small MPs and even nanoplastics (smaller than 1 µm (Choi et al., 2020; Gigault et al., 2018)) can cause apoptosis, increased reactive oxygen species (ROS), and reduced microbiome diversity (Al-Jaibachi et al., 2019; Brandon et al., 2018; Peng, Xu, et al., 2023; Richard et al., 2024; Wang et al., 2021). In planarians, ingestion of MP beads and fibres caused increased levels of apoptosis, but this effect was only evident in individuals consuming the smallest size of beads (10 µm) (Gambino et al., 2020). Snails (*Cantareus aspersus)* feeding on the smallest size class of MPs (50-100 µm) suffered from oxidative stress, while those ingesting larger plastics did not (Colpaert et al., 2022). Similarly, in an aquatic chironomid (*Chironomus tepperi)*, only the smallest class of MPs (1-4 µm) caused a reduction in survival, growth, and body size (Ziajahromi et al., 2018). Overall, the ingestion of smaller particles appears to have more severe impacts than the ingestion of larger MPs’ (Agathokleous et al., 2021; Sucharitakul et al., 2024); but such effects likely scale and interact with body size.

Plastics change during passage through the insect gut, and these changes can have lasting negative effects on the environment. Insects can physically break down plastics into smaller pieces resulting in the release of even smaller plastics in the animals that consume them, and the excretion of smaller plastics into the environment (Cassone et al., 2020; Peng, Sun, et al., 2023; Peng, Xu, et al., 2023; Ritchie et al., 2024). Insects with larger heads are known to generate more force through mandibular power due to the increased size of their mouthparts (Bernays, 1986; Edel et al., 2024; Hochuli, 1996; Laiolo et al., 2021). Some insect orders, like Orthoptera and some Diptera, also increase the forces applied to ingested food by increasing the mineral density in their mouthparts during development and reducing abrasion from ingesting hard food (Laiolo et al., 2021; Lehnert et al., 2022; Winkler et al., 2024). As a result, insects should be able to apply more force to their food, and break down harder foods as they develop (Laiolo et al., 2021; Lehnert et al., 2022). The same process may also allow for the increased breakdown of MPs during ingestion. Due to terrestrial insects being important food sources for many insectivores, insect interactions with MPs can drive trophic transfer of plastics and bioaccumulation (Cui et al., 2022; Huerta Lwanga et al., 2017; J. Zhang et al., 2024). However, the extent to which insects ingest, modify, and transfer modified plastics to predators has not been thoroughly studied.

Generalist crickets, common in agricultural settings, will ingest plastics in both field and laboratory settings (McColville et al., 2023; Ritchie et al., 2024). In the lab, crickets (*Gryllodes sigillatus*) will ingest large quantities of plastic when given no choice but to eat contaminated food (Fudlosid et al., 2022; Ritchie et al., 2024). This phenomenon can also be found in nature; the fall field cricket, *Gryllus pennsylvanicus*, collected from fields treated with biosolids, had a wide range of MPs of different sizes within their digestive tracts (McColville et al., 2023). Crickets (*G. sigillatus*) can also degrade ingested plastics to smaller particle sizes as adults and appear to suffer few outward effects from ingestion of ∼100 µm MPs during lifelong feeding (Fudlosid et al., 2022; Ritchie et al., 2024). These findings suggest that terrestrial insects, including crickets, could drive transfer of MPs to higher trophic levels, and body size and particle size likely interact to impact whether insects ingest plastics and transfer them. The model *G. sigillatus* can be used to understand how body size, particle size, and biodegradation within an insect can all independently and, when combined, interactively affect the probability of trophic transfer within natural settings.

Given the limited knowledge of how body size impacts plastic ingestion and excretion, we examined the consumption and passage of microplastics in a model cricket (*G. sigillatus*) to address two hypotheses: First, we hypothesized that mouthpart size would limit the size of plastic particles that an insect can or would ingest because particles are consumed whole. We predicted that crickets would ingest smaller plastics very early in development and that larger plastics would only become available for ingestion later in later developmental stages when the cricket was bigger. Second, we hypothesized that body size and microplastic size would impact the degree to which plastics were broken down in the cricket digestive system. We predicted less plastic breakdown would occur in the early stages of cricket development, when forces generated are weaker, and greater plastic breakdown would occur later in development in concert with increased body size.

## Methods

### Cricket Rearing

Cricket rearing was conducted following previously described methods (Ritchie et al., 2023). Briefly, *G. sigillatus* eggs came from the colony at Carleton University initiated from a colony from Entomo Farms, an industrial partner that raises crickets for food and feed. Eggs were raised in a greenhouse at ∼32°C and 40% RH with a 14:10 L:D light cycle. Once the pinheads (1^st^ instar cricket nymphs) hatched, approximately 500 were placed in each of three bins (60 cm × 40 cm × 31 cm) with cardboard egg trays provided for shelter and *ad libitum* water and access to food (made of corn, soybean, herring, and hog meal mixture). Crickets were left in these bins for seven weeks under the same conditions for the mouth size measurements (see below).

Crickets for body measurements and frass collection were individually housed in 3 ¼ oz plastic condiment cups at one week old with water provided in 0.65 mL microcentrifuge tubes with dental cotton plugs, and diet provided *ad libitum* in microcentrifuge tube lids under the same L:D and humidity. Crickets in this experiment were fed the standard diet with MPs mixed into the feed until they reached seven weeks of age. Typically, adulthood is reached within six weeks at 32°C and at this time body size and mass largely plateau (Fudlosid et al., 2022; Kong et al., 2024). The microplastics mixed with the feed were low-density fluorescent blue polyethylene microspheres 1.13 g/cc (Cospheric) ranging from 38-500 µm with clearly defined size thresholds of 38-45, 75-90, 150-180, 250-300, and 425-500 µm (henceforward referred to as 38, 75, 150, 250, 425 MPs).

Crickets for the *diet choice* experiment came directly from Entomo farms at the age of three weeks and were housed in bins (as listed above) until they reached adulthood. The *diet choice* experiment used the 38 MPs and another size class of 90-106 µm (henceforward known as 90 MPs) as small and medium-sized MP to assess whether crickets respond differently to different size classes of plastics.

### Diet Choice Experiment

After reaching adulthood (∼ 6 weeks), individual male and female crickets were exposed to a pairwise combination of two of the following three diets: 1) a control diet, made of the base feed with no microplastics, 2) a 38 MP diet, made of the base feed with 0.5% w/w microplastics, and 3) 90 MP diet, made of the base feed with 0.5% w/w microplastics. Crickets were pseudorandomly assigned to one of the three pairwise treatments: 1) control and 38 MP diet, 2) control diet and 90 MP diet, and 3) 38 MP diet and 90 MP diet.

Each feeding dish (14 mm wide polyethylene push-in cap) containing food was separated into individual feeding trays on either end of an opaque black rectangular container (19 cm x 12.7 cm x 5 cm, #PC838BLACK, polypropylene, CBS Distributing) with clear lids modified with wire mesh. Water was provided in microcentrifuge tubes and a dental cotton covering. Feeding dishes were dried in an oven at 30°C for ∼48 h and then weighed before and after removal from the container. The three different diet choice scenarios were presented individually to ten males and ten females (60 crickets total). Crickets were weighed before being individually placed inside and left to feed for nine days, with water replacement every two days and food replacement every three days. Every three days, the bins were rotated throughout the incubator and diet positions were swapped each time they were weighed. Crickets were also weighed every three days during the food swaps, and the remaining food was returned to a 30-degree oven to dry for approximately seven days before final weight values were collected. The remaining food from each cricket enclosure was separated from any visible frass and weighed to calculate the amount of food ingested from each feed dish. These values were then used to determine which food was preferentially ingested in those three days. After the nine-day trial, crickets were removed from the containers and re-weighed.

### Effects of Plastic on Body Size

One week old crickets were pseudorandomly assigned to groups of 50 from the three bins of ∼500 crickets. The 50 crickets were housed individually and exposed to the control diet or the same base diet containing 1% w/w of microplastics of a single size class (38, 75, 150, 250, or 425 MPs) for the duration of the experiment. Cricket frass was collected every seven days when the plastic cup was cleaned, and the frass was frozen to measure the plastic quantity and the degree of biodegradation. The collection happened within 24 h of body measurements on the same individual crickets. Body size was quantified as both width and length of the head and thorax, with measurements taken as described previously (Fudlosid et al., 2022; Kasdorf et al., 2025). Water and diet for the respective cricket were refilled every 2-3 days until seven weeks for all groups with respective diets. The frass was stored at −20°C for future digestion. Survival of the crickets from each group was also tracked throughout the experiment. A total of 40 out of the 300 crickets died due to accidents during handling or measurements. These 40 crickets were removed from all analyses.

### Plastic Degradation in the Gut

The digestion method for frass was repeated from Ritchie et al., 2023. The frozen frass from each cricket was removed from the freezer and allowed to thaw. Once thawed, the frass pellets from each cricket were placed in a 0.6 mL storage tube to be digested with 30 µL of 10% KOH and incubated for two days at 60°C. After two days, the samples were vortexed and filtered on a 1 µm pore size glass fibre filter (Sigma, APFB04700) on top of a vacuum pump assembly (Sigma, Z290408-1EA). The MPs were filtered using a hand pump (Fisher, S12932) attached to the vacuum pump assembly. After filtering, the filter paper with the MPs was moved to a petri dish and photographed under a dissection microscope (Zeiss, Stemi 508 with an axiocam 105 colour).

Following methods from Ritchie et al. (2023), isolated fluorescent plastic particles in the images were analyzed using ImageJ (FIJI version 2.3.051; Java 1.8.0_172 [64-bit]). The analysis involved separating the colour channels of each image and removing the red channel to reduce noise. The blue and green channels were then re-stacked and used to quantify particles using ImageJ’s watershed and binary settings. Output from this allowed for measuring the surface area µm^2^ of each particle and the roundness (4*area/(π*major_axis^2) of particles present within each sample. Methods of isolation and measurements were conducted on non-digested food (master mix) of each bead size to allow separation between whole and broken-down beads.

### Mouth Size

Accurate measurements of mouthpart size require sacrificing animals. Therefore, to examine how ingested particle sizes scale relative to the mouth and head of the cricket, mouthparts of group-reared crickets were measured following dissection in a separate set of animals. The *mouth size* group was left on the control diet for all seven weeks and reared like the control crickets (see above). Ten crickets were pseudo-randomly selected from bins at one-week intervals during development to measure their mandibles, maxilla, labium, and labrum. Other body parts (the head, thorax, abdomen, and leg) were measured on the same animals. After photos of the ventral and dorsal side of the cricket were taken, the head was removed for dissection. Dissection was done using an insect pin inserted through the ventral side of the head in a Sylgard-lined petri dish (World Precision Instruments, SYLG184). Then, the maxilla from both sides was removed, followed by removing the labium with dissection forceps. The mandibles were then removed with a micro scalpel and dissection forceps, followed by the removal of the labrum. All six mouthparts were then arranged to be photographed and measured (Figure 2).

Mouth and body size measurements were completed in Fiji (ImageJ) v2.3.051. This was done for head width, thorax width, thorax length, abdomen width, abdomen length, femur length, mandibles, maxilla, labium, and labrum. Mouthparts were measured by previously defined width and length measurements (Bertram et al., 2021; Judge & Bonanno, 2008; Krenn et al., 2016). *Effects of plastic on body size* were measured using Napari (v0.4.17). Key points were placed at the widest points of the head (for head width), the thorax (for thorax width), and at the top and bottom of the thorax (for thorax length). Distances between these key points were extracted using a custom Python script (v3.9), yielding head width, thorax width, and thorax length. Sex, when possible, was determined through visual inspection for all experiments.

### Data Analysis

Data was cleaned for impossibly large bead sizes particles (which result from artifacts from large particles close together on the filter, Table S1) for each MP group by removing all MPs larger than two standard deviations above the mean of the control beads (Ritchie et al., 2023). The proportion of removed observations for each size case was calculated (Table S1). An informal review of these images confirmed that the removed particles were too close together, leading to implausibly large particle areas as described previously (Ritchie et al., 2024). All plotting and statistical analyses were conducted using R version 4.1.2 (R core team, 2023). Unless otherwise stated, the lmer model was used with a type III Anova in all experiments, and cricket ID was treated as a random effect in all models. The ratio within the behaviour data was calculated as the amount of plastic-containing food ingested divided by the total amount of food ingested. The choice between the plastic sizes 100 µm over 38 µm was the ratio of 38 µm food ingested over the total amount of ingested food. A ratio of whole beads to broken-down particles was calculated from imaged particles in the frass. Particles were only considered a breakdown product if the area and roundness of the beads were more than two standard deviations below the mean value of the control beads for each size of microplastics fed to the crickets (Table S1, Figure 4).

## Results

From our choice experiment, we compared the amount of food ingested for each diet - one containing plastic (38 or 90) and one uncontaminated. Values greater than 0.5 indicate a greater proportion of ingestion of plastic food (Figure 1A, B) or a preference for the diet containing smaller MPs (38 µm) when both diets contained plastic (Figure 1C). The sex of the cricket did not influence its diet choice when both plastic diet options were available (Sex; F_1, 28_ = 0.31, p = 0.58). No cricket absolutely avoided any of the diets presented to them during each sampling period (Figure 1). When given a choice between uncontaminated and plastic-contaminated food (38 or 90 µm), crickets first had no preference for uncontaminated food but then significantly shifted towards the plastic-contaminated food after nine days (Figure 1A, B) (Sample period; F_2, 43_ = 3.64, *p* = 0.033). The size of the plastic particles (38 or 90) presented with clean food did not affect preference over time (Bead size; F_1, 47_ = 2.55, p = 0.121). Crickets did not prefer 38 or 90-sized plastics when only plastic-contaminated diets were given (Sample period; F_2, 25_ = 3.0, *p* = 0.068).

**Figure 1.**
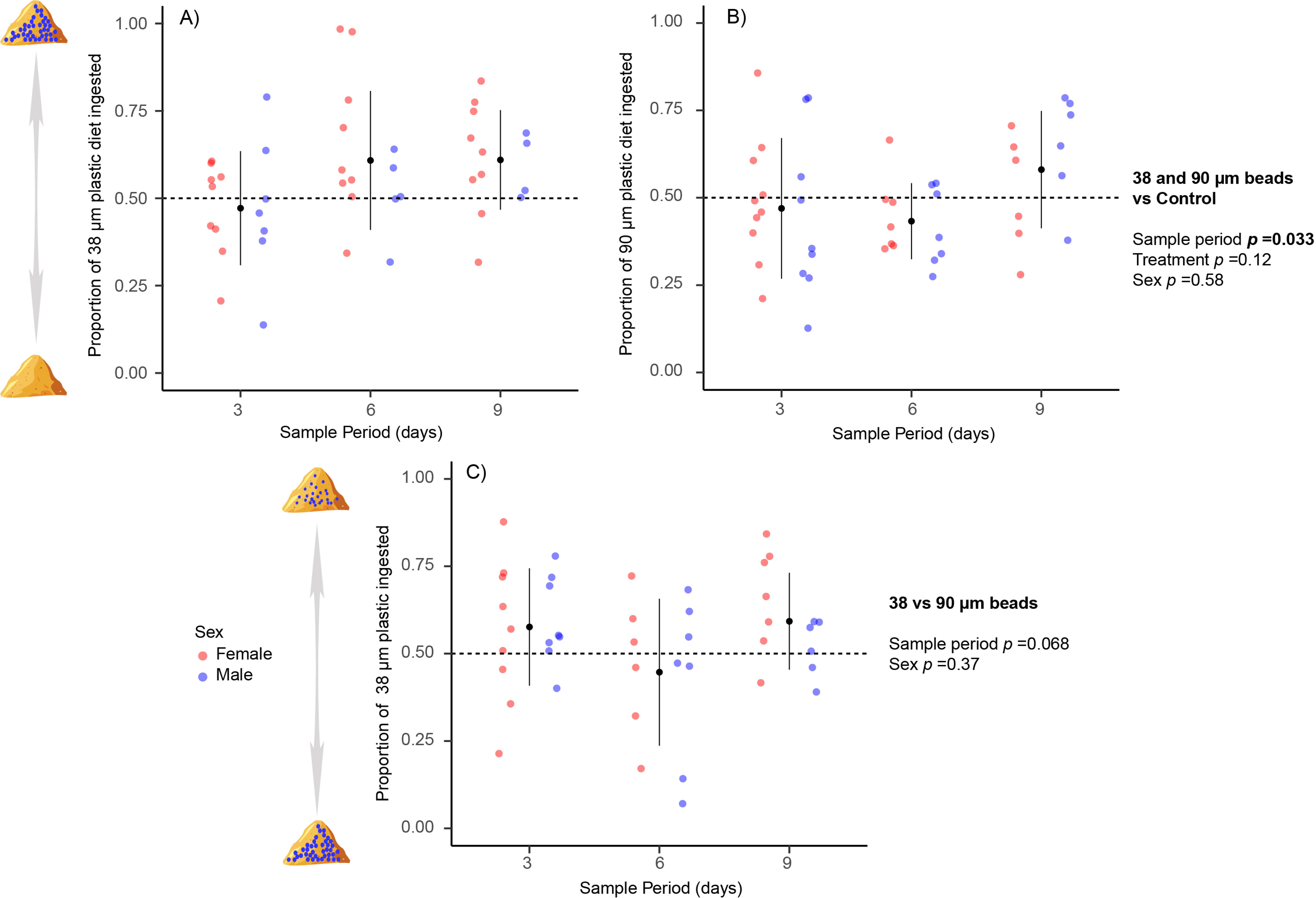
The proportion of feed consumed by individual male (blue) and female (red) crickets (*G. sigillatus*) during a choice experiment where either uncontaminated or microplastic-contaminated food was presented (A and B) or where two contaminated diets were provided containing two size classes of MPs (C). Sampling took place over nine days, with diet positions swapped during each sampling period. Male and female crickets show slight but significant changes in preference toward contaminated food as exposure time increases. Crickets showed no preference in the size of the plastic beads in the feed. Individual coloured points represent individual crickets, and black points represent group means ± SD.

To better understand how body size interacts with particle size to influence the probability of plastic ingestion, we quantified the ingestion of several sizes of MP beads over the entire developmental period of the cricket. We were particularly interested in how plastic ingestion scales with mouth size, so we first related mouthpart size to common indices of body size using principal component analysis (PCA) (Figure 2A). From this, we confirmed that the labium (lower jaw, Figure 2B) of the crickets correlated best with non-sacrificial body measurements that were also collected in our plastic ingestion and breakdown experiment. Specifically, the labium correlated strongly and positively (r^2^ = 0.98) with the cricket’s head and thorax width (Figure 2C). We therefore opted to use the head width as a predictor of labium size in the ingested particle size experiment. We created a linear model from our mouth size experiment to predict labium size in all crickets fed a plastic diet (Figure 2D).

**Figure 2.**
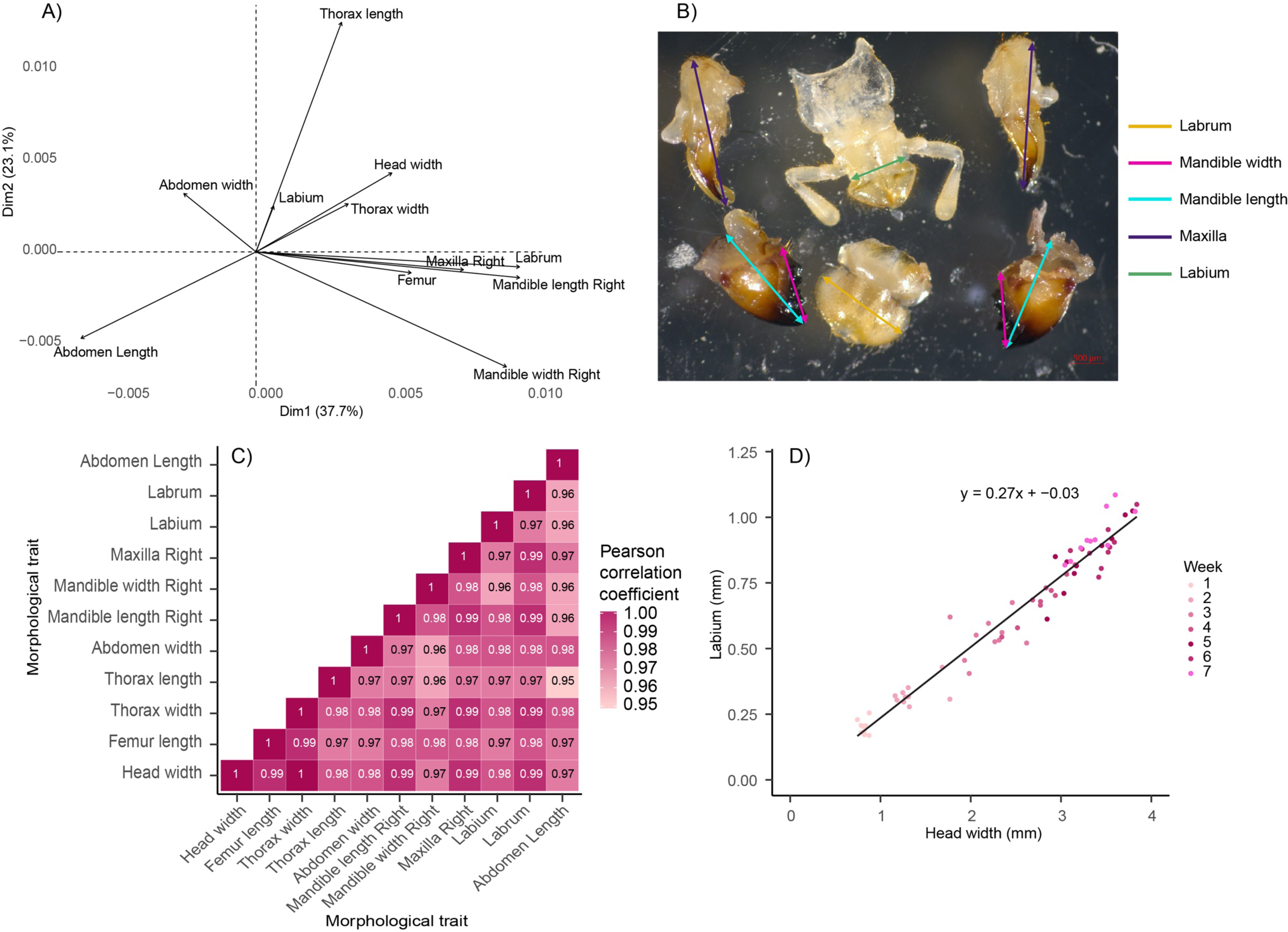
Relationships between multiple measures of mouthpart size and body size in *G. sigillatus* throughout development. PCA analysis of body and mouth size indices (A). Example image of mouthparts after dissection (B). Between-index Pearson’s correlation coefficients (C). Linear correlation of male and female crickets between the head width and their labium with linear equation (D). Head width and thorax width (non-sacrificial measurements) strongly correlate with the size of the mouthparts and can, therefore, be used to approximate mouth size during development. Week refers to the week of cricket development during which crickets were sacrificed, and individual points in panel D represent individual crickets.

Using the head width as a predictor of mouth size, we sought to understand how patterns of MP ingestion and breakdown changed as mouth size increased. We first measured growth over all weeks of both male and female crickets to see whether any size classes of MPs affected growth (Figure S2). Regardless of the MPs present in the diet and sex, we found no effects of plastic ingestion on body size. This was the case regardless of whether we considered head width (male: χ²(5) = 2.21, *p* = 0.81; female: χ²(5) = 0.69, *p* = 0.98 (Figure 3 A and B)), thorax width (male: χ²(5) = 0.49, *p* = 0.99; female: χ²(5) = 0.30, *p* = 0.99 (Figure S2)), or thorax length (male: χ²(5) = 0.71, *p* = 0.98; female: χ²(5) = 0.25, *p* = 0.99 (Figure S2)).

**Figure 3.**
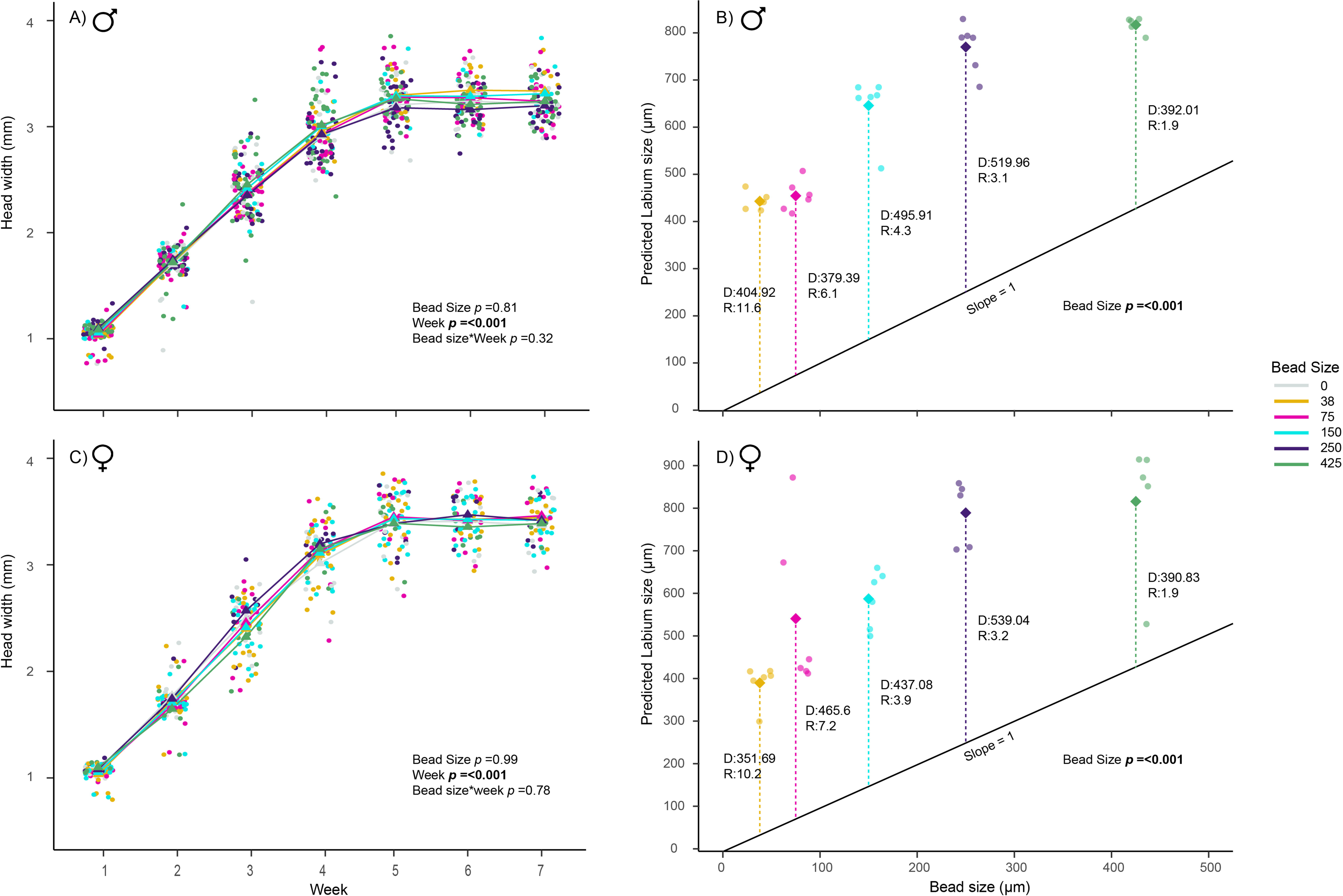
Mouth size predicts the likelihood of plastic particle ingestion across ontogeny in crickets (*G. sigillatus*). Lifelong feeding on different-sized MPs does not affect changes in the growth of male and female crickets (A and C). Head width correlates well with labium size (Figure 2D), and was used to predict labium size from their head width. As labium size increased for both male and female crickets, breakdown products of MPs from larger beads appear in the frass, indicating that crickets can ingest them (B and D).

We next explored when plastics were first excreted (as evidence of ingestion) across all bead sizes using a pseudorandomly selected subset of six males and six females from each treatment group. We found that the timing of the first excretion of plastics was heavily dependent on the bead size presented in the food. Plastics were only found in the frass of crickets presented with larger plastics when they themselves were larger. The size of the plastic present in the frass was limited by the size of the labium in males (Bead size; F_1, 27_ = 114, p < 0.001) and females (Bead size; F_1, 26_ = 33, p < 0.001) crickets. At the first examination of the frass (2 weeks of age), the two smallest bead sizes, 38 and 75, were present within the frass of both male and female crickets. However, the remaining 150, 250, and 425 sizes were absent from the frass until crickets had grown and the labium exceeded ∼500 µm. Labium size at first ingestion scaled approximately linearly with the size of the bead ingested, such that the labium was consistently between ∼350 and ∼540 µm larger than the bead before evidence of ingestion was found in cricket frass. We found that the ratio of MP size to mean labium size decreased as the labium and MP size increased (represented by R in Figures 3 B,D).

The digestive system of crickets dramatically degrades ingested plastics, but this has previously only been demonstrated using a single-size class of beads. To better understand how body size and ingested particle size interact to influence MP breakdown, we analyzed the area and roundness of egested particles relative to control beads of each MP size. We found that the control beads were very consistent in size, as expected, but we didn’t know what proportion of whole beads vs fragments were present within the frass samples of the crickets. We aimed to compare the MPs found within the male frass from all weeks vs. the control beads to understand how much crickets were breaking down each size class of bead (Figure 4). Using two standard deviations of the area and roundness from the control beads, we defined the limits of particles we would consider a whole bead still present within the frass. All particles outside these defined limits were considered fragments created by the crickets. All bead sizes were found to have a relatively normal distribution of roundness and area (Figure 4); however, control samples of the 38 and 75 beads (Figure 4 A, B) did contain some smaller plastics. Regardless, all control beads were highly clustered, indicating that plastics mixed into the feed were almost exclusively whole beads within the expected size class. It was also clear that the median area and roundness of beads in the frass were lower than the controls (e.g. Figure 4 E; Area of controls equals 192461 µm^2^ vs excreted equaling 125 µm^2^). This indicates that once these crickets ingest these plastics, they cause a large amount of change.

**Figure 4.**
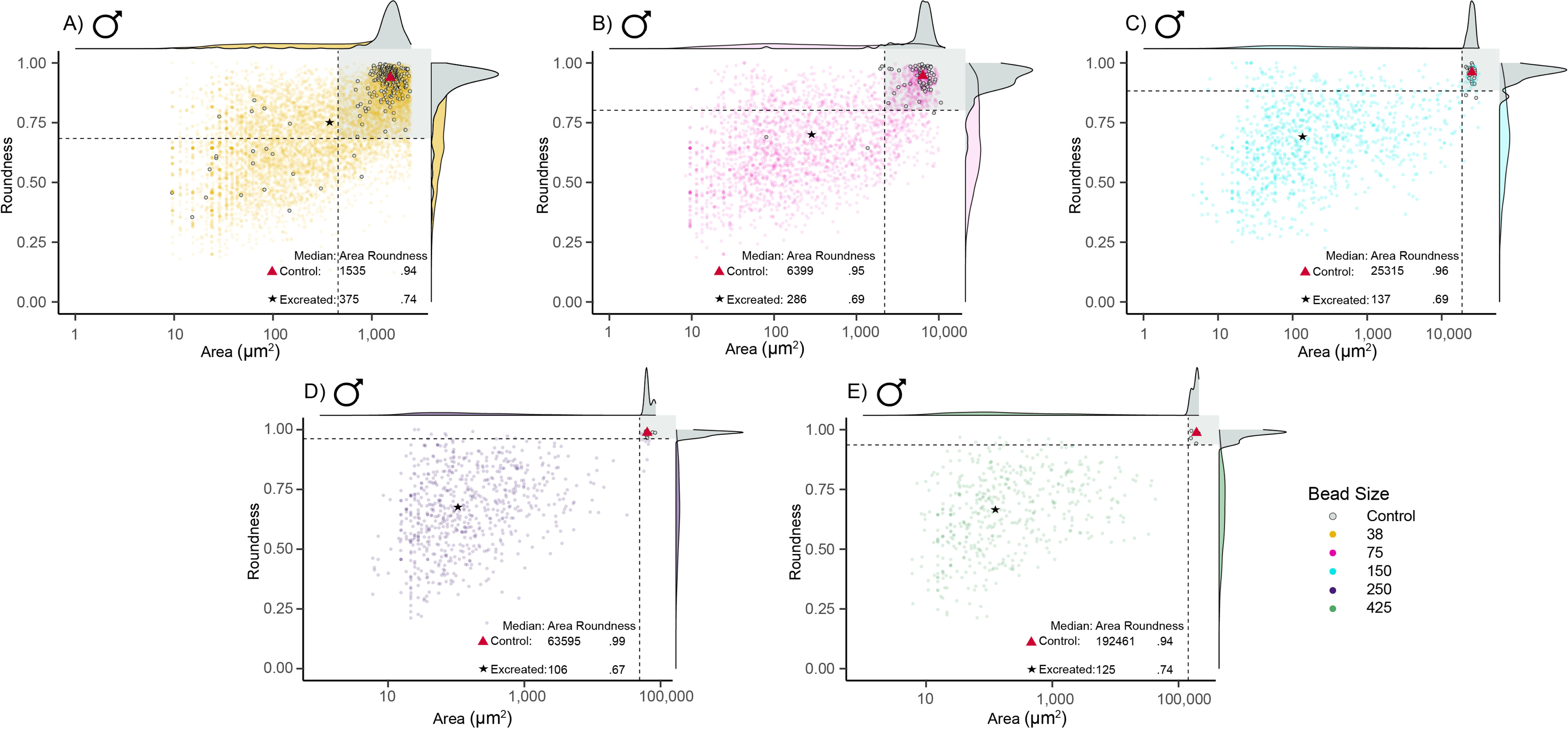
When ingested, larger size classes of MPs are more fragmented in the digestive system of male *G. sigillatus*. Compiled figures showing the defining points for a whole bead (gray background) vs fragment (white background) in the frass of male crickets. Control beads (grey) are shown within each panel for each size of MP (A-E). All bead fragments found within all weeks of all male crickets used in the study are shown (coloured circles). Dashed lines represent two standard deviations lower than the mean area and roundness of control beads. Density plots along axes show the distributions of particle area and roundness.

Particle size clearly impacted the degree to which breakdown occurred within the cricket gut. To better understand how sex, development stage, and bead size impacted the magnitude of fragmentation of MPs within the digestive system, we calculated a ratio of whole vs. fragmented particles in the frass of male and female crickets during each week of development. Sex, ingested bead size, and developmental week (here used as a proxy for body size) all interacted to influence the total amount of plastic excreted in the frass (Sex*Bead size*Week; χ²(15) = 264, *p* = < 0.001). Because of this interaction, we visually separated these results by sex (Figure 5B, C). For both sexes, the smallest beads (38 and 75 µm) were typically found in a consistent ratio of 1 (meaning equal parts unfragmented beads and breakdown products), except for females in week two, when even less breakdown was observed. As mouth size increased, the 150 class of beads became available for consumption; the breakdown of this size class tended to become less common as crickets grew, particularly in females. The 250 and 425 size classes of beads also became available for consumption as the labium size further increased. For these larger MPs, the degree of breakdown significantly differed between males and females (Sex; χ²(1) = 216, *p* = < 0.001). Over time, this ratio increased for females fed the 150 and 250 size class beads, indicating they excreted a greater proportion of whole beads as they continued to grow. By contrast, the males only showed such an increase in the ratio for the 150 beads and this trend was more variable. For males, whole 250 size class beads were only noted as present once. Regardless of sex, crickets ingested the largest bead size (425), but no whole 425 beads were ever found in the frass. Females grew faster and larger than males and ingested the 250 and 425 class beads before the males started to ingest these size classes.

**Figure 5.**
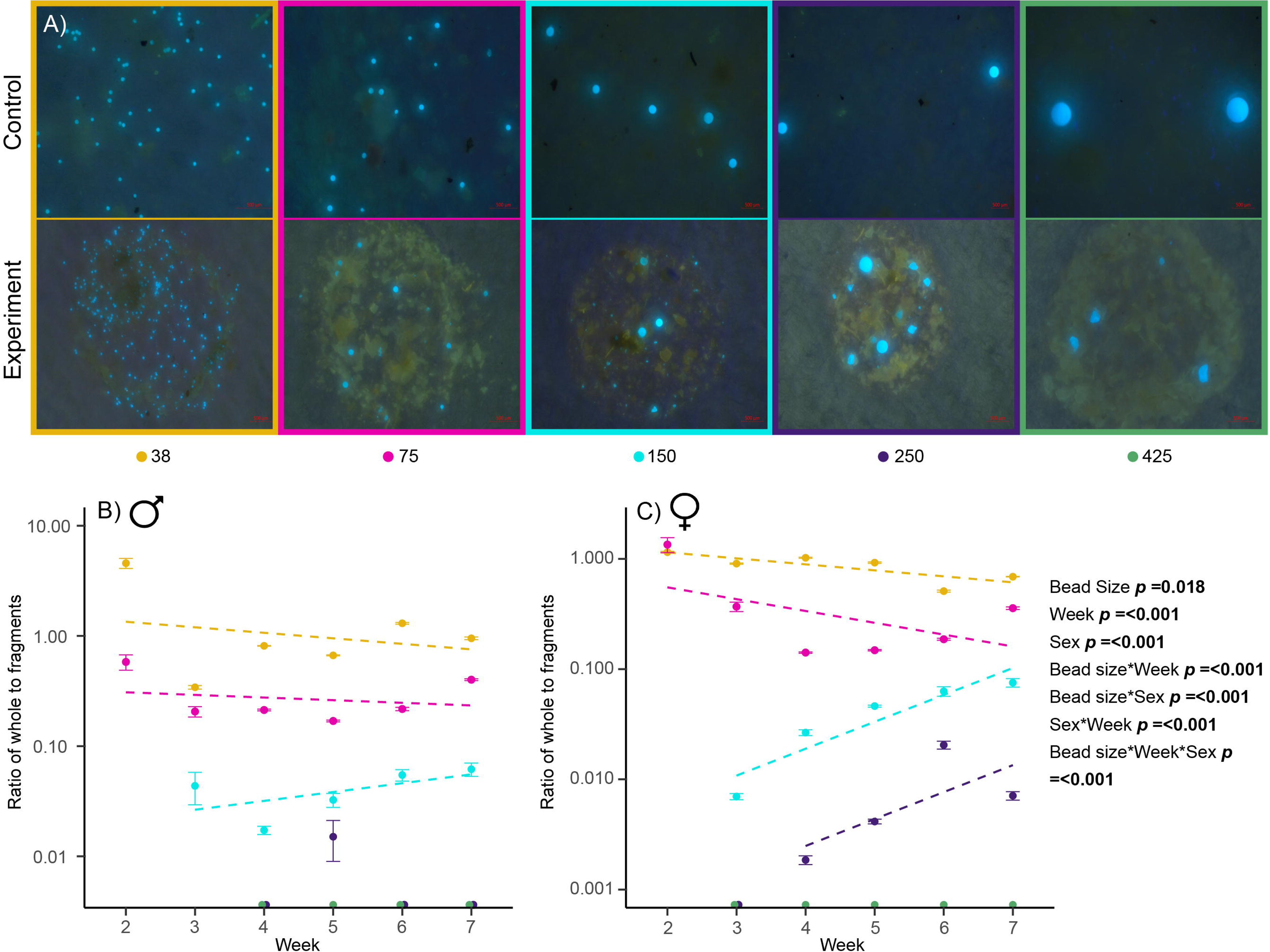
Microplastic fragmentation depends on complex interactions between particle size, body size, and sex in crickets (*G. sigillatus*). Despite some fragmentation, smaller beads are excreted whole throughout all of ontogeny, intermediate size beads are fragmented less as the body size of the cricket increases, and large beads are always fragmented once ingested. Images show examples of how different bead sizes change during transit through the gut as the crickets develop, all images are female crickets as they appeared in the frass (A). Significant differences exist between the amount of breakdown of the different MPs as the crickets grow over time (B, C).

## Discussion

Developing a clear understanding of how invertebrates interact with plastics is critical to understanding the full impacts of plastics on ecosystems, and thus informing policies that include considerations of these impacts. Because plastics vary in size, polymer, colour, and chemical additives, building this knowledge represents a significant challenge that necessitates both top-down and bottom-up approaches. Here, we sought to use a model insect to specifically understand how cricket body size interacts with plastic particle size to modulate ingestion and transformation. Given that MPs are defined within a 5000-fold range of sizes (1 µm to 5 mm), we expected both the probability of ingestion and the degree of transformation to change through ontogeny in this cricket species.

In a lab setting, crickets don’t seem to avoid or seek contaminated food sources regardless of particle size. When exposed to an environmentally relevant concentration, crickets initially slightly preferred food without MPs, but this preference declined over time regardless of sex and the size of MP present. Since contaminated environments typically have a wide range of MPs sizes present, crickets are unlikely to consume plastics of the same size throughout their lives. Changes in feeding behaviour in response to MPs have been documented in flies and bees when they were given no choice but to ingest MPs along with food (Balzani et al., 2022; Ranjan et al., 2024). Recent evidence suggests that ants initially avoid MP-containing food but lose this preference over time (Le Hen et al., 2024). Wild Orthopterans, such as crickets, have plastic in their bodies, suggesting that insects may not avoid plastic in contaminated natural environments (Lu et al., 2020; McColville et al., 2023). Together, the evidence suggests that a wide taxonomic group of insects cannot distinguish plastics from food, a particularly troubling conclusion given the growing prevalence of plastic in natural environments.

We found strong support for our first hypothesis that crickets are limited to consuming MPs that are small enough to pass through their labium (lower jaw). Only the smallest size classes of MPs (38 and 75) were ingested early in development (at two weeks of age); larger MPs (150, 250, 425) were only ingested later in development after the labium had increased in size. Our findings are comparable to those observed in *C. riparius* larvae when fed different size classes of MPs ranging from 1-90 µm (Scherer et al., 2017). Plastic ingestion began only when the head capsule width of *C. riparius* reached a size sufficient to ingest MPs. In our study and in Scherer et al. (2017), both insect species were initially limited to ingesting small MPs when their head capsule or labium was small. As both insect species grew, their larger heads enabled them to grasp and ingest progressively larger MPs, with ingestion continuing throughout their development. Together, these findings suggest that small insect species, and/or insects early in development, have access to all plastic particles smaller than their mouth size. Smaller plastics are more detrimental (Brandon et al., 2018; Choi et al., 2020; Richard et al., 2024), and our research indicates that most terrestrial insects could ingest these harmful smaller plastics. Our findings contribute to our fundamental understanding of insect MP ingestion in two crucial ways. First, they suggest that larger insects are likely to ingest a more comprehensive range of plastics, as once ingestion starts for a particle size, it continues throughout development. Secondly, because crickets did not ingest the large MPs in their food (until their mouth size increased), we argue that crickets are unlikely to ingest MPs larger than they can ingest whole and are therefore unlikely to remove smaller pieces of plastic from larger objects (macroplastics; >5 mm).

When ingested, larger particles were more extensively broken down by the cricket digestive system, while smaller particles were more likely to be excreted whole throughout ontogeny. This finding supports our second hypothesis that changes in cricket body size would impact the breakdown of particles, but the direction of our prediction was incorrect. We found all sizes of MPs had the highest production of fragments to whole particles when first ingestion became possible. An alternative explanation for this pattern is that larger particles were trapped in the digestive system, so whole beads could not pass. We argue that this is unlikely since we saw no difference in survival or growth, regardless of the particle size ingested (Figure S3). This finding suggests that smaller whole particles allow more effective passage through the digestive tract and decrease the opportunity to become NPs (smaller than 1 micron). Hemimetabolous insects (with incomplete metamorphosis) are likely to contribute to the increased creation of MPs and NPs once ingested. Holometabolous insects (with complete metamorphosis) like mealworms and silkworms can break down MPs during their larval phase, but breakdown products excreted by these species do not appear to reach the NP level (Peng, Sun et al., 2023). Developmental differences in plastic degradation could be related to whether insects use mechanical or enzymatic mechanisms to break down the microplastics. Regardless, both mechanisms would facilitate the transfer of plastics or their metabolic products through the food chain. If the patterns of plastic breakdown observed here hold for other terrestrial invertebrates beyond crickets, our findings could be important for waste management decisions; body size ranges of the dominant animals within an environment could dictate the degree to which, and the rate at which, plastic fragmentation occurs.

We found that once crickets can ingest a particle whole, they can and will continue to ingest that particle size for the rest of their lives. Here, the labium was consistently 350-540 µm wider than a plastic particle before we observed evidence of ingestion (Figure 3), suggesting a consistent “clearance” in the digestive tract was needed before crickets could ingest the plastic particle. By contrast, as black soldier fly larvae develop, they ingest plastic particles equal in size to their mouth (Lievens et al., 2023). Insects are known for having extremely diverse mouthparts that relate to their food sources, and among-species differences in plastic ingestion behaviour may be attributed to variations in mouthpart or digestive tract anatomy. Like crickets, black soldier fly larvae are chewing insects. Black soldier fly larvae have a joined mandibular maxillary unit as their primary feeding apparatus (Armstrong, 2001; Bruno et al., 2020), whereas crickets have separate mandibles and maxillae capable of food manipulation (Bertram et al., 2021; Krenn et al., 2016). To more broadly identify the factors governing relationships between body or mouthpart size and the probability of plastic ingestion, we argue that effort should be put toward testing these relationships across a diverse group of species representative of different feeding styles. We further expect patterns of plastic breakdown occurring through invertebrate interactions will shift seasonally as age classes and dominant species change.

The sex of a cricket determines the amount of fragmentation that occurs, specifically with larger particles that become available later in life. Because female crickets are generally larger than males, they had more whole beads present (observed for the 150, 250, and 425 MP sizes) in their frass as they developed (Fudlosid et al., 2022; Kong et al., 2024). The smallest beads used here (38 and 75) were broken down to a consistent degree across ontogeny for both sexes. This does not support our prediction that smaller crickets would be less able to generate forces needed to break down the plastic. Female crickets started ingesting larger MPs earlier in their development than male crickets. Additionally, female crickets did not break down the ingested plastics into smaller pieces as extensively as males did before excreting them. These differences in ingestion and plastic breakdown may be attributed to the fact that female crickets grow larger than males as they develop, which allows females to ingest larger MPs earlier. This sex difference may also affect how plastics are processed in cricket bodies. The sizes of 150 and 250 MPs showed higher fragmentation initially, but as the female crickets increased in body size, more whole beads were found. A possibility for the increased presence of whole beads could be due to the size of the cricket gut increasing with age. Future studies could investigate if morphological changes in the gut allow for the passage of large particles. No whole 425-sized MPs were ever present in the frass despite clear evidence of plastic ingestion, indicating it is beyond the maximum size that can pass through the digestive system. In our study, both sexes could break down these large beads (425 and 250), but males mostly did not pass 250 beads whole and instead broke them down in the gut. This indicates a potentially higher threshold for females of ingestible sizes than the males. Future studies should consider sex differences in plastic availability as sexual dimorphism in body size, relative head capsule size, and relative mouth part size found in many species (Bertram et al., 2011; Judge & Bonanno, 2008).

### Conclusions

Overall, we report that a model cricket species does not avoid a plastic-contaminated food source and may even develop a slight preference toward it. Crickets readily consumed and degraded plastics of various sizes, but only once they themselves were large enough to eat the MP particles whole. Crickets can break down polyethylene throughout their whole life, but the degree to which MP particles are broken down after consumption depends on the initial particle size and cricket body size, with larger particles generally being more heavily degraded. Based on these findings, we argue that the biodiversity and abundance of invertebrates could strongly impact the degree to which microplastics move through our natural environments.

## Acknowledgments

We thank Entomo Farms for supplying cricket eggs during the project’s development. Marshall Ritchie also thanks Rebecca Dean for many late-night conversations on this work during the data collection. The authors would also like to thank Sophie Kasdorf and Émile Vadboncoeur for assisting in colony maintenance and cricket care.

## Author Contributions

**Marshall Ritchie:** Conceptualization, Methodology, Validation, Formal analysis, Investigation, Data curation, Writing - Original Draft, Writing - Review & Editing, Visualization. **Emily McCoville:** Conceptualization, Methodology, Investigation, Data curation. **Jennie Mills:** Investigation, Data curation**. Jennifer Provencher:** Conceptualization, Methodology, Writing - Review & Editing, Supervision, Funding acquisition. **Susan Bertram:** Formal analysis, Methodology, Writing - Review & Editing, Supervision. **Heath MacMillan:** Conceptualization, Methodology, Writing - Original Draft, Writing - Review & Editing, Visualization, Supervision, Funding acquisition.

## Data availability

All data is provided as a supplementary file for review, and the same file will be included as supplementary material should the manuscript be accepted for publication.

