## Supplemental for "The disadvantage of having a big mouth: the relationship between insect body size and microplastic ingestion"

Table S1: Control bead measurements and values used for data curation

| Bead size | Mean Area ($\mu m^2$) | Mean Roundness | Mean and roundness - SD*2 | Highest particle size ($\mu m^2$) | Ratio of particles removed (%) |
|---|---|---|---|---|---|
| 38 | 1467 | 0.903 | 455, 0.684 | 2479 | 1487/ 20088 (7.4) |
| 75 | 6162 | 0.929 | 2199, 0.802 | 10124 | 47/ 6723 (0.70) |
| 150 | 25777 | 0.953 | 18715, 0.883 | 32840 | 15/ 2869 (.52) |
| 250 | 67127 | 0.982 | 49257, 0.961 | 84997 | 1/ 2688 (.037) |
| 425 | 184501 | 0.977 | 141968, 0.936 | 227033 | 0/1211 (0) |

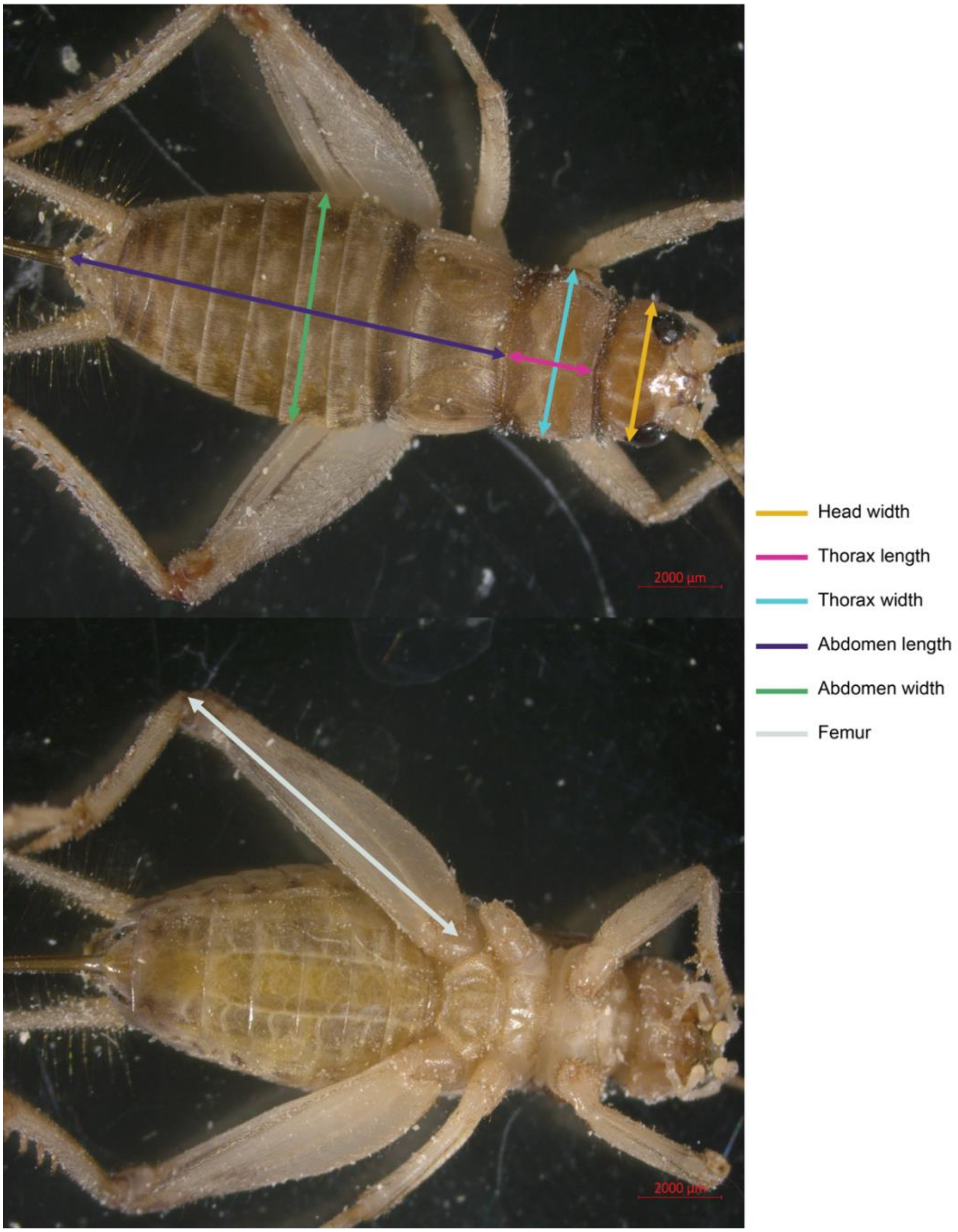


Figure S1: Morphological measurements of crickets that did not require sacrificing. Arrows dictate the placement to be measured using AI and manual measurements.

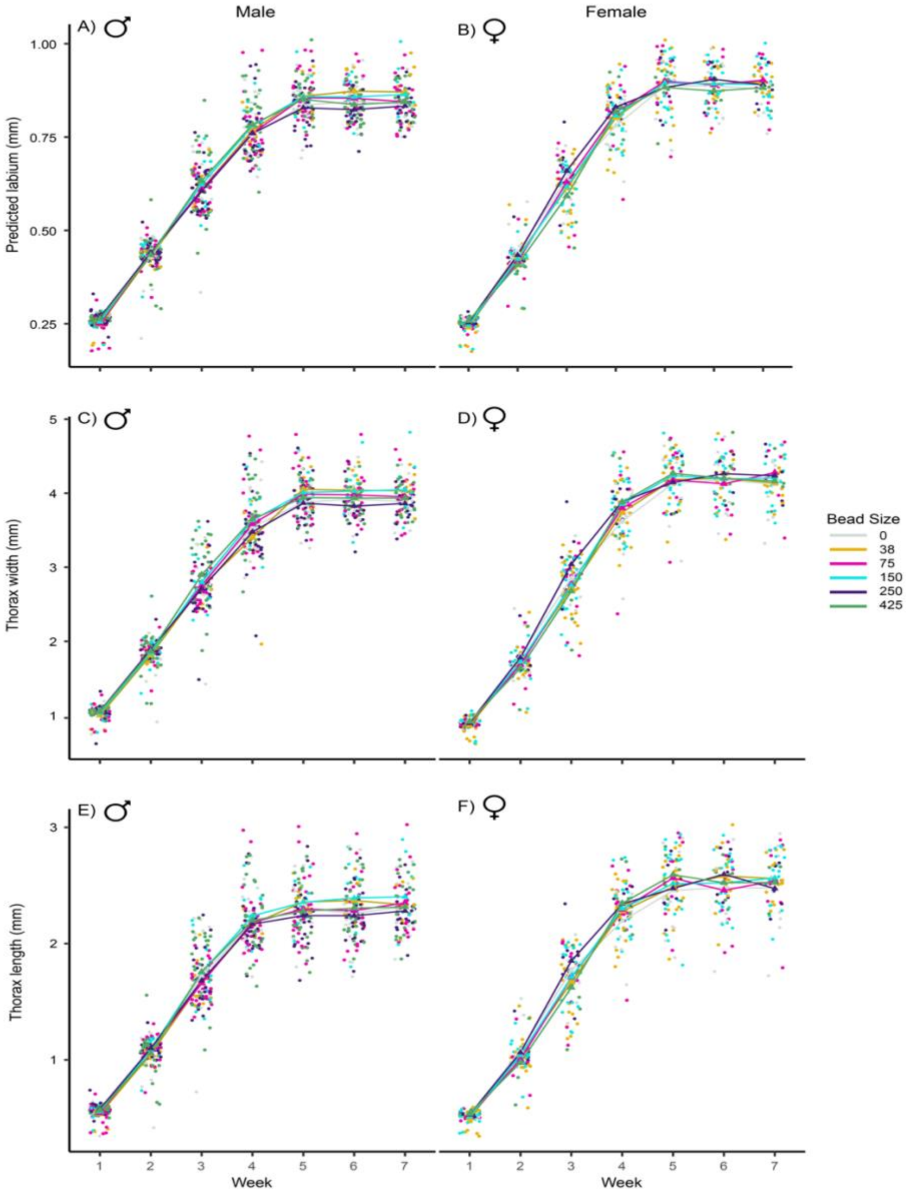


Figure S2: Male and female cricket body metrics over lifelong feeding of MP beads. Predicted labium of all crickets in the experiment throughout their development (A and B). Thorax width of crickets throughout their development (C and D). Thorax length of crickets as they aged throughout their development (E and F).

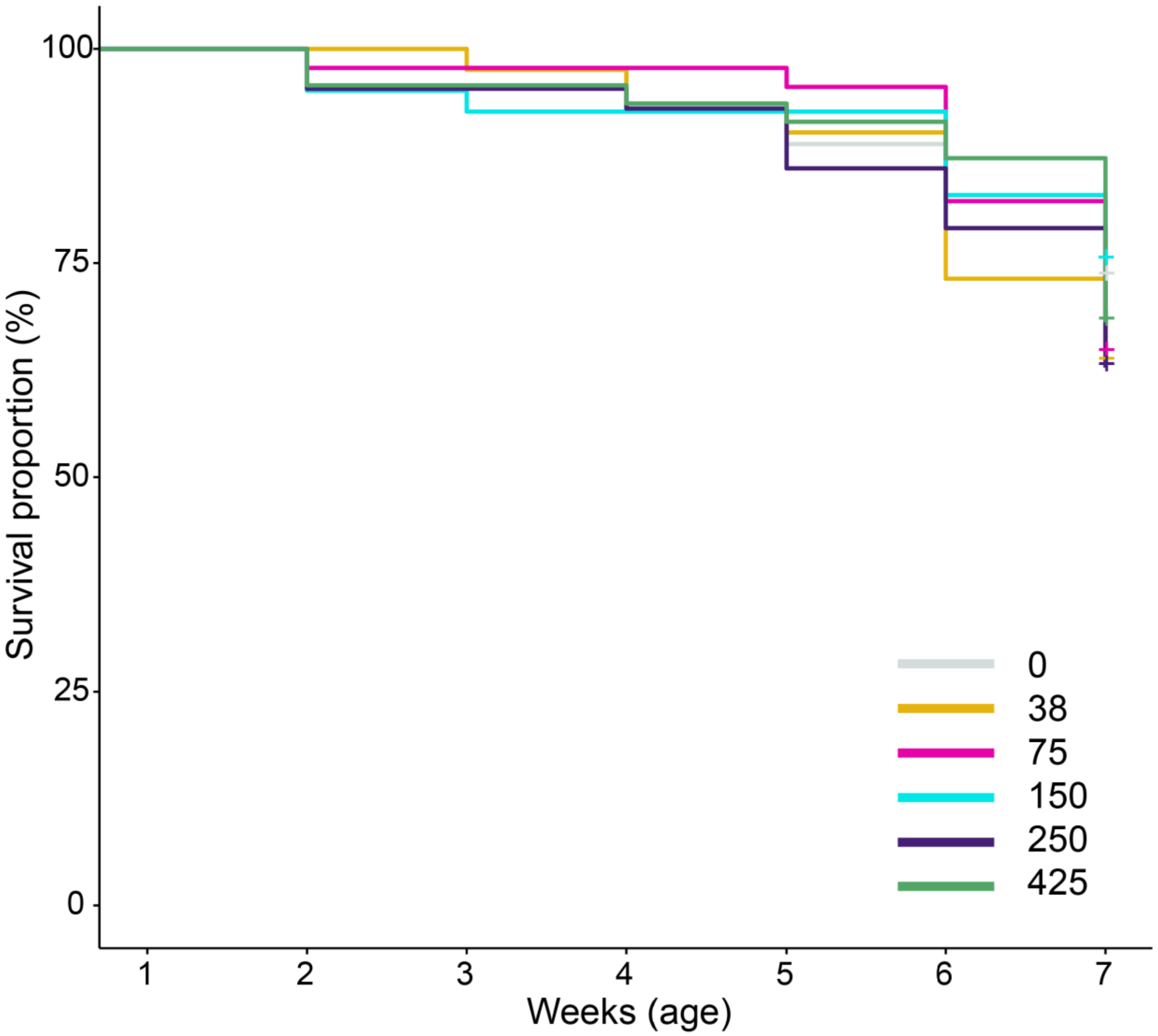


Figure S3: Survival analysis of crickets fed different bead sizes throughout their development. No difference was found for any of the treatments for the crickets. A log-rank test was conducted to compare the survival distributions among different bead sizes (38, 75, 150, 250, and 425 µm). The survival distributions were not significantly different between the groups, $\chi^2(5) = 2.6$, $p = 0.8$. This suggests that bead size does not have a statistically significant effect on the survival time of the crickets.
